# ERC 2.0 - evolutionary rate covariation update improves inference of functional interactions across large phylogenies

**DOI:** 10.1101/2025.02.24.639970

**Authors:** Jordan Little, Guillermo Hoffmann Meyer, Aakash Grover, Alex Michael Francette, Raghavendran Partha, Karen M. Arndt, Martin Smith, Nathan Clark, Maria Chikina

## Abstract

Evolutionary Rate Covariation (ERC) is an established comparative genomics method that identifies sets of genes sharing patterns of sequence evolution, which suggests shared function. Whereas many functional predictions of ERC have been empirically validated, its predictive power has hitherto been limited by its inability to tackle the large numbers of species in contemporary comparative genomics datasets. This study introduces ERC2.0, an enhanced methodology for studying ERC across phylogenies with hundreds of species and tens of thousands of genes. ERC2.0 improves upon previous iterations of ERC in algorithm speed, normalizing for heteroskedasticity, and normalizing correlations via Fisher transformations. These improvements have resulted in greater statistical power to predict biological function. In exemplar yeast and mammalian datasets, we demonstrate that the predictive power of ERC2.0 is improved relative to the previous method, ERC1.0, and that further improvements are obtained by using larger yeast and mammalian phylogenies. We attribute the improvements to both the larger datasets and improved rate normalization. We demonstrate that ERC2.0 has high predictive accuracy for known annotations and can predict the functions of genes in non-model systems. Our findings underscore the potential for ERC2.0 to be used as a single-pass computational tool in candidate gene screening and functional predictions.

## Introduction

The -omics era has greatly increased the number of sequenced genomes (Hotaling et al. 2021). With this influx of information, network and systems biologists have developed approaches to study gene and protein interactions from a broader perspective and at a much larger scale. Networks of such interactions can be reconstructed based on gene co-expression, semantic similarity, and other experimental methods. Such networks have become commonly used to select disease gene candidates (Chen et al. 2009; Paredes-Sánchez et al. 2015; Sun et al. 2010) and to predict protein function (Saha et al. 2019; Sharan et al. 2007; Xiong et al. 2014; Zhu et al. 2010). However, there are still limitations. These methods require many contextual details to be understood, such as the environment from which the organism/cells were collected, the phenotypes being expressed, and the developmental stage from which the information is captured. The requirement for this *a priori* knowledge limits many studies to studying only model systems. By studying protein interactions through a phylogenetic lens, we can remove the need for these details and incorporate non-model species into the analyses. Understanding that gene expression, physical interactions, and epistatic interactions all potentially respond to the selection pressures on a gene, we can study patterns of evolution across species to find genes and proteins that have shared functions and processes. In this study, we propose an improved evolutionary method to build systems biology-level networks that allow for analysis of both model and non-model species.

Evolutionary selective pressures act on every species across different developmental stages, environments, and phenotypes. Thus, we can treat each species as its own experiment conducted across millions of years. Adding more observations in a traditional bench experiment increases the power to detect patterns. Similarly, adding more species to a phylogeny increases the statistical power to detect evolutionary patterns. These patterns can be used to identify genes that have been subjected to shared selective pressures. Since genes that participate in a function together (*i.e.*, co-functional genes) experience many of the same selective pressures (Goh et al. 2000; Pazos and Valencia 2001; Clark et al. 2012), changes in those pressures are expected to change their relative evolutionary rates (RER) in parallel. Thus, by identifying genes with correlated changes in branch-specific RERs, functional relationships between genes can be inferred.

Evolutionary rate covariation (ERC) is a measure that quantifies the correlation of the RERs of a pair of genes across all branches in a phylogeny (Figure 1A). ERC is not to be confused with simple correlations between the average rates of genes; rather, ERC examines how rates change between species and whether those changes correlate with other genes. A high ERC correlation value is an evolutionary signature of co-function: as is demonstrated by the RERs of SMC5 and SMC6, two proteins that are known to form a complex (r = 0.78, Figure 1B). While SMC5 and SMC6 show high ERC, the expectation is that most gene pairs are not co-functional and so would not be acted upon by shared selective pressures. To illustrate, proteins that are not functionally related tend to have correlation values closer to zero, as seen when SMC5 (r = 0.15) or SMC6 (r = 0.06) is compared to a non-related gene PEX6 (Figure 1A,B).

**Figure 1:**
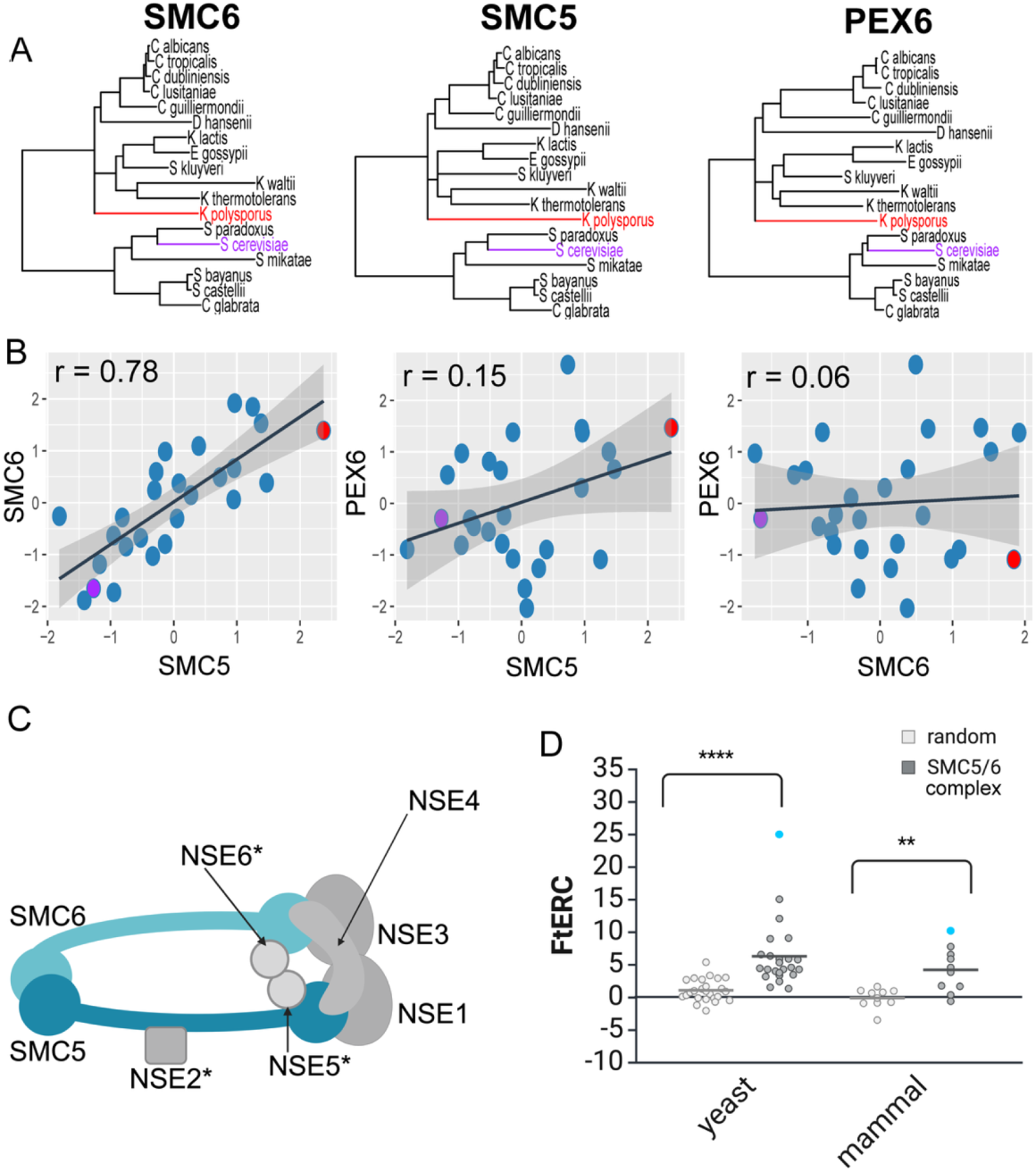
SMC5/6 complex shows significantly elevated ERC in both yeast and mammal phylogenies. A. Gene trees showing the branch lengths for SMC6, SMC5, and PEX6 respectively across 18 yeast species. Representative species, *S. cerevisiae* and *K. polysporus*, are highlighted in purple and red respectively, in all three trees. B. Scatter plots comparing the relative evolutionary rates for SMC6 x SMC5, SMC5 x PEX6 and SMC6 x PEX6 using the same 18 yeast species as shown in A with the representative species, *S.cerevisiae* and *K. polysporus*, shown in a purple and red dot, respectively. A linear regression line with 95% confidence interval is shown in black and gray respectively. The Pearson correlation for each comparison is shown in the top left corner of the plot. C. Cartoon schematic of the SMC5/6 complex (* indicates a yeast-specific complex member). D. Fisher transformed ERC (FtERC) values for all pairs within the SMC5/6 complex for a yeast dataset of 343 species (left, gray) and mammal dataset of 120 species (right, gray) with size matched random gene pair samples for each dataset (white). The datapoint with the highest FtERC in both datasets is between SMC5 and SMC6 (blue dot). Both datasets had significantly higher ERC (**p< 0.001, ****p<0.00001) than the sample of randomly selected gene pairs from the entire genome. Panels C and D created in BioRender. Little, J. (2025) https://BioRender.com/o07i564.

In addition to the high ERC between SMC5 and SMC6, ERC values between all pairs of proteins in the SMC5/6 complex are positive and elevated in a dataset using 343 yeast species (Shen et al. 2018) and for almost all pairs in a dataset using 120 mammal species (Hecker and Hiller 2020) (Figure 1C, D). These observations reflect the co-functional relationship of these proteins, as they have experienced the same changes in selective pressures over evolutionary time. In general, the ERC scores are higher for the yeast dataset than for the mammals, likely due to the greater amount of evolutionary divergence among the yeast species. These results demonstrate how ERC matrices are useful as a mineable resource for evolutionary screening for new functional relationships in various taxa.

Calculating ERC for each pair of orthologous genes across a phylogeny allows for the generation of an evolutionarily derived co-functional map of the cell, which in turn can be used to predict functional relationships and interactions (Goh et al. 2000; Pazos and Valencia 2001; Clark et al. 2012; Yosef et al. 2009). ERC has been calculated on a wide variety of taxonomic groups, including mammals (Priedigkeit et al. 2015; Kowalczyk et al. 2021), fungi (Clark et al. 2012; Little et al. 2024; Clark et al. 2013; Steenwyk et al. 2022), *Drosophila* (Findlay et al. 2014; Raza et al. 2019), and plants (Forsythe et al. 2021).

In principle, high ERC values allow the detection of any type of functional relationships, whether underpinned by physical interactions, such as protein complexes (Little et al. 2024), or non-physical interactions, such as enzymatic pathways (Findlay et al. 2014; Brunette et al. 2019; Raza et al. 2019). ERC has been used to successfully identify novel interactors in protein pathways that were then experimentally validated (Findlay et al. 2014; Raza et al. 2019; Brunette et al. 2019). It has also been used to screen for candidate genes in disease networks (Priedigkeit et al. 2015), identify relaxation of constraint in meiotic proteins in clonally reproducing yeast (Clark et al. 2013) and determine the intracellular localization of poorly characterized zinc transporters (Kowalczyk et al. 2021). The success of ERC as a screening method demonstrates the efficacy and usefulness of the method across phyla and biological scopes. However, the ERC methodology used in the studies above was only performed on a maximum of 63 species.

Here, we introduce a new open source software package ERC2.0. This software makes it possible, for the first time, to calculate ERC across hundreds of species and tens of thousands of genes. Whereas previous methods used Pearson correlations and branch lengths not corrected for heteroskedasticity, ERC2.0 makes improvements to employ Fisher transformation and a new branch length normalization step, each of which we demonstrate to improve statistical power. Our case studies on datasets of yeast and mammal species (supplementary file1, supplementary file 2) show the improved predictive power of ERC2.0 relative to our previous implementation, ERC1.0 (Clark et al. 2012).

## Results

### Fisher transformation normalizes correlations and reduces variance across diverse branch counts

Previous iterations of ERC have been used to validate protein interactions (Clark et al. 2012), identify candidate genes within pathways (Findlay et al. 2014), and build gene-based disease networks (Priedigkeit et al. 2015) using the Pearson correlation coefficient. However, in larger datasets, such as a dataset of 120 mammals (Hecker and Hiller 2020), the number of branches contributing to the correlation can vary when a gene is not present in every species. This creates two issues. First, the correlations cannot be compared across datasets because of the discrepancy in the number of data points contributing to each score. Second, with fewer branches, the distribution of correlation coefficients exhibits a greater variance (Figure 2A).

**Figure 2:**
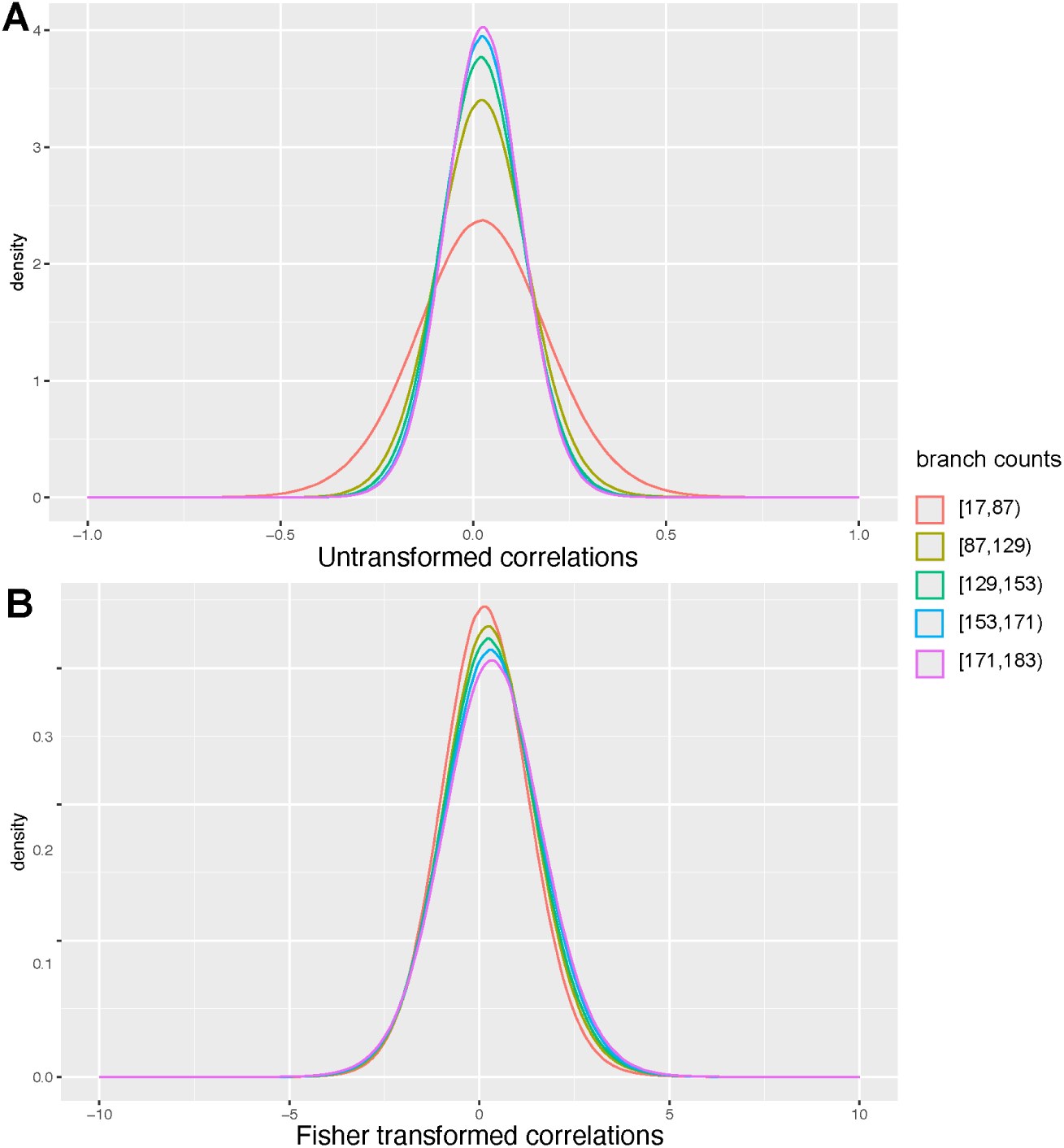
Fisher transformation normalizes variance across gene pairs with different branch counts. A. Untransformed Pearson correlations for gene pairs calculated on 120 mammals. Gene pairs were sorted into equally sized bins based on the number of branches that contributed to the ERC score. B. Fisher transformed correlations for gene pairs calculated on 120 mammals. Gene pairs were sorted into equally sized bins based on the number of branches that contributed to the ERC score.

To normalize the correlation coefficients for the number of branches that went into the calculation, we introduce a Fisher transformation (Fisher 1915) to ERC values using the following equation:

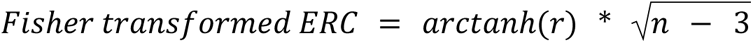

Where *r* is the correlation coefficient, and *n* is the number of branches. Over a population of correlation coefficients, this transformation yields a normal distribution from [-∞, ∞] with stable variance (Figure 2B). The null expectation of no correlation remains at zero.

This transformation allows scores to be meaningfully compared across taxa. Using Fisher-transformed values results in a more consistent variance and higher predictive power than untransformed correlation coefficients (<u>Figure S1</u>).

### Faster: ERC2.0 allows for computationally tractable calculations of hundreds of species

Increasing the numbers of species and genes permits investigating novel interactions but requires a tractable compute time. ERC2.0 uses a new data structure to speed the computation of all correlations between all genes. A major challenge in computing all pairwise correlations is that it requires pruning each gene’s tree so that their species set matches; most genes are missing several species due to evolutionary or technical reasons. For this reason, ERC2.0 stores all trees in a format that includes all possible subtrees. Together with a rapid indexing system and efficient C-based tree operations in the TreeTools package (Martin R. Smith 2019), the retrieval time for matching trees for each gene pair was greatly decreased, which makes datasets of much larger size accessible (<u>Table S1</u>). While ERC1.0 takes over 2 hours to calculate ERCs between 500 genes for 343 species on 10 CPUs (<u>Table S1</u>), ERC2.0 with this new methodology completes the same task in 4 minutes.

### Removal of heteroskedasticity in branch rates improves power

An additional improvement is that RERs on each gene tree branch are now calculated using a method that removes heteroskedasticity from the resulting rates, as was developed for the RERconverge package (Partha et al. 2019). Because these rates no longer have a variance that scales with their magnitudes, they are better suited in theory for application in statistical tests, such as linear correlations (ERCs). In practice, we measured a clear increase in power when branch RERs were corrected for heteroskedasticity (ERC2.0) compared to uncorrected (ERC1.0), as seen by the improvement in area under the ROC curve (Figure 3, 0.652 to 0.581, respectively).

**Figure 3:**
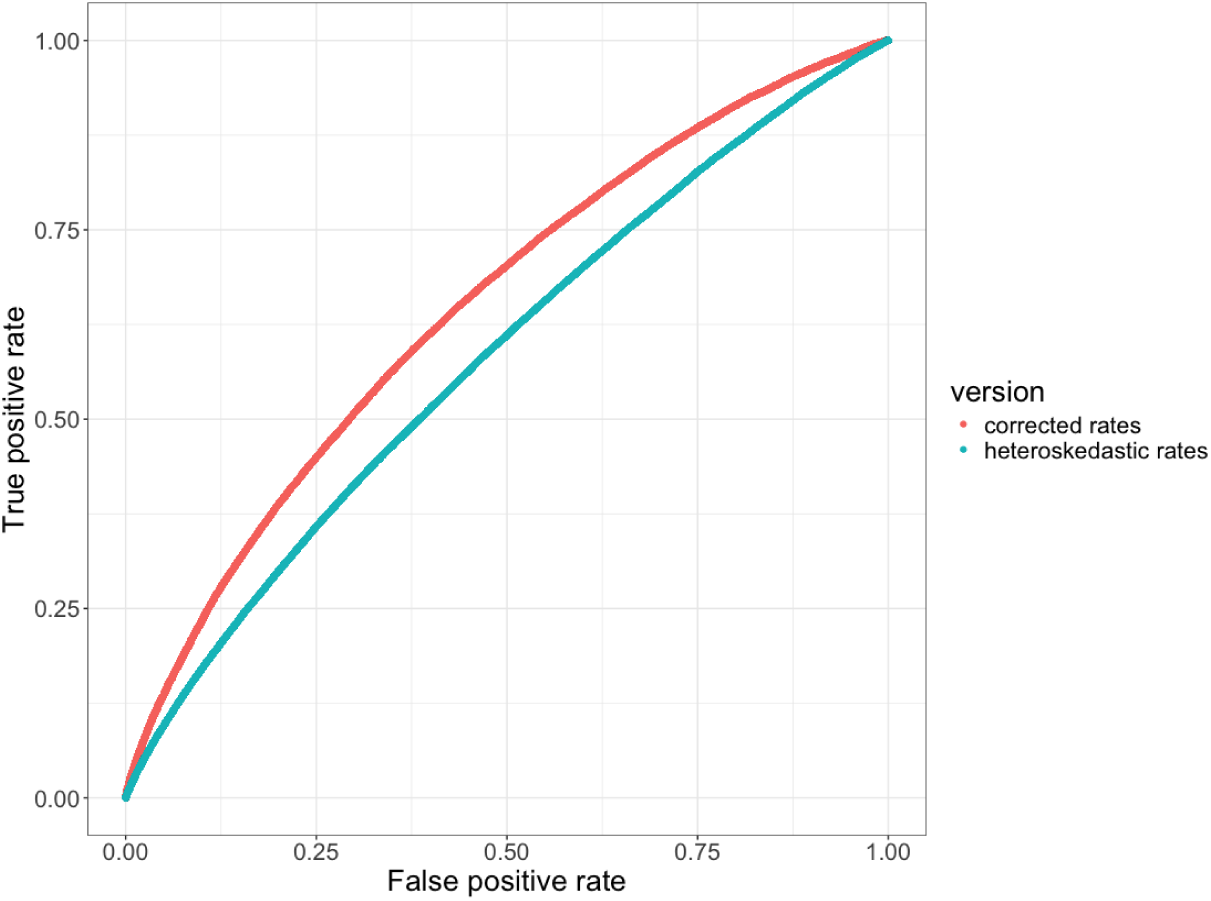
Removal of heteroskedasticity improves ERC predictive power. Receiver Operator Characteristic (ROC) curves contrasting true positive and false positive rates demonstrate higher power in ERC2.0 values calculated using corrected rates (red curve, AUC = 0.652) compared to ERC1.0 values using uncorrected, heteroskedastic rates (blue curve, AUC = 0.581). The true positive set are interactions from STRING with combined scores > 700. ERC scores were calculated on a phylogeny of 18 yeast species.

### Better: ERC2.0 outperforms previous ERC methodology and improves with larger datasets

The faster methodology can calculate ERC scores for tens of thousands of genes across hundreds of species. To evaluate the increase in power when more species are added, we expanded the analysis of a previous ERC study that used only 18 species of yeast (Clark et al. 2013). We hypothesized that increased species counts would improve the biological relevance of ERC scores, therefore, predictive power.

To test this we used logistic regression modeling to test how well ERC can be used to predict which genes belong to a given functional annotation (<u>Methods</u>). Logistic regression modeling allows us to test the practicality of using ERC scores to predict gene function for binary outputs where a gene either belongs to an annotation or does not. The logistic regression model then uses the ERC values between all genes to select which genes or “features” are most important for accurately predicting the membership in a particular functional annotation. We used 10-fold cross-validation with an elastic net penalty and the area under the ROC curve (ROC-AUC) as the evaluation measurement to optimize the model. This resulted in a set of ROC curves across different regularization parameters that allows us to choose the model with the best fit for an annotation term.

We began by querying the broad TORC1 signaling pathway, which has elevated ERC (Figure 4A). For this pathway, ERC2.0 calculated on 343 yeast species produced a ROC-AUC of 0.988, meaning it had very little error in predicting members of that pathway (Figure 4B). We then looked at more annotations to test how well ERC can predict pathways and processes globally. We used two datasets for yeast and mammals found in yeastEnrichr (Kuleshov et al. 2016) or Enrichr (Kuleshov et al. 2016), respectively. We again used ROC-AUC to quantify the performance of each ERC dataset to predict two different sets of annotations (Figure 4C-F). In yeast we used KEGG pathways (Kanehisa 2019) and GO-Biological Processes (GO-BP) (Ashburner et al. 2000; The Gene Ontology Consortium et al. 2023). For both sets, ERC2.0 run on 343 yeast species had a significantly higher average ROC-AUC than both ERC2.0 run on 18 species and ERC1.0 on 18 species. While the average ROC-AUC for ERC2.0 run on 18 species was not significantly higher than for ERC1.0 run on 18 species, it was still higher for GO-BP (0.72 vs 0.71 respectively) and KEGG (0.77 vs 0.73). Even more promising, the average ROC-AUC for ERC2.0 run on 343 yeast species is greater than 0.8 for both GO-BP and KEGG, indicating that the ERC2.0 dataset has a high potential for predicting new genes within the annotated functional terms.

**Figure 4:**
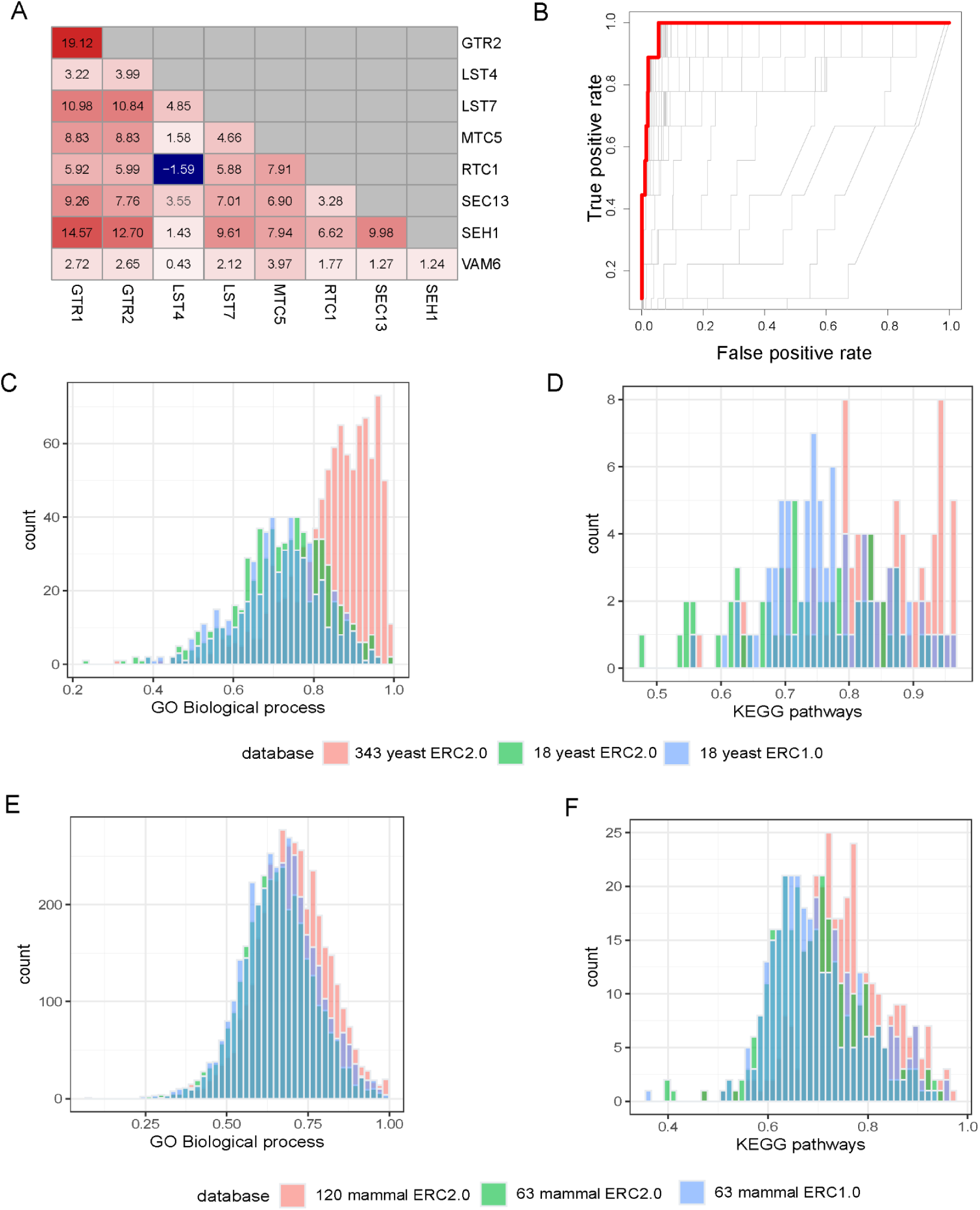
ERC2.0 improves predictive potential for GO Biological Processes and KEGG pathways. A. Pairwise yeast ERC scores for members of the Biological Processes pathway, ‘Positive regulation of TORC1’. Red colors indicate a higher FtERC, blue indicates a lower FtERC. B. ROC curves for each model generated with cv.glmnet for ‘positive regulation of TORC1’. The red line indicates the model with the highest ROC-AUC of 0.988. C-D. cv.glmnet validation of GO biological processes (C, n=545), and KEGG pathways (D, n=49) using ERC2.0 on 343 yeast species (red), ERC2.0 on 18 yeast (green) and ERC1.0 on 18 yeast (blue). There is a significant difference in means between 343 yeast ERC2.0 and both 18 yeast ERC2.0 and ERC1.0 for both GO Biological processes (p<2.2e-16, p<2.2e-16, respectively) and KEGG pathways (p<2.2e-16, p = 1.089e-08, respectively). E-F. cv.glment cv.glmnet validation of GO biological processes(E, n=5807) and KEGG pathways(F, n=320) using ERC2.0 on 120 mammal species (red), ERC2.0 on 63 mammal species (green) and ERC1.0 on 63 mammal species (blue). There is a significant difference in means between 120 mammal ERC2.0 and both 63 mammal ERC2.0 and ERC1.0 for both GO Biological processes (p<2.2e-16, p = 1.147e-05, respectively) and KEGG pathways (p=1.259e-15, p=1.486e-12). Each count is the highest AUC (s=lambda.min) predicted by glmnet.cv.

Since the difference in ROC-AUC between the original 18 yeast dataset and the current 343 yeast dataset is large, we also performed ROC analysis in intermediate datasets by subsetting the 343 species dataset in increments of 49 species (Figure S2).

We found a plateau as the number of species reached ∼100. We hypothesize that the initial increase is due to clade-specific interactions as species are added and that the plateau is due to the saturation of annotated genes in the dataset.

Both GO-BP and KEGG were also tested on the three iterations of mammal ERC. Similar to the yeast datasets, the larger 120 mammal dataset run with ERC2.0 had a significantly higher mean AUC than either of the 63 mammal datasets run with ERC2.0 or ERC1.0. While the 120 mammal ERC2.0 dataset showed a lower average ROC-AUC for both datasets than yeast with the average AUCs at 0.70 and 0.77 for GO-BP and KEGG, respectively, there are still terms that have a ROC-AUC above 0.9. This shows that there are annotation terms that ERC2.0 is especially primed to predict in mammals.

We also compared the predictive power of ERC to the current literature base as captured in STRING (Szklarczyk et al. 2019). We used lambda.1se AUCs for the same two yeast annotation datasets as mentioned above to compare models trained using ERC values to models trained using STRING confidence scores (<u>Figure S3</u>). We limited the ERC analysis to only genes that are also present in STRING. The AUCs for the STRING-trained models were greater than the ERC AUCs for 1147/1166 of the annotations, with a mean AUC of 0.95.

We next asked whether the combination of ERC and STRING scores would improve even the STRING model performance (<u>Figure S3</u>). Combining them improved the AUC of 1363 annotation terms compared to STRING alone, indicating net improvement upon adding ERC. The greatest improvement was for the GO Biological Process term “peptide transport (GO:0015833)”, which improved from a STRING AUC of 0.742 and ERC AUC of 0.723 to a combined AUC of 0.914. While ERC is unable to outperform the entirety of the STRING dataset, the improvement of some annotation terms, as well as the relatively high average AUCs for ERC alone, show that there is novel information being captured by this tool. Furthermore, we envision that ERC2.0 will provide rapid and highly accurate functional prediction when applied to non-model organisms lacking the extensive experimental datasets found in yeasts and mammals, and it can be done using sequence information alone for a relatively minuscule cost.

### ERC2.0 networks best predict clusters of organism-level interactions in mammals compared to cellular processes in yeast

Given the comparatively low ROC-AUCs for the mammalian datasets, we investigated the overall structure of the mammal ERC interaction network compared to the yeast ERC network to determine the types of functional interactions best captured in each. We first subsetted each dataset to the top 1% of FtERC values to capture the strongest set of gene-gene interactions. We then used a Markov chain clustering (MCL) algorithm in Cytoscape (Shannon et al. 2003) to create clusters of genes based on similar ERC profiles.

The resulting yeast network consisted of 257 clusters, with cluster 1 containing 3346 of the 4181 genes (<u>Figure S4</u>, supplementary file 1). We performed GO: Biological process enrichment tests (Thomas et al. 2022) on the 12 clusters with greater than 10 members (Figure 5). This allows us to determine what types of processes within the cell/organism are shared by the genes within a cluster. Given the density of cluster 1, we performed another round of MCL clustering on only that cluster which broke it into 7 sub-groups. These sub-groups were largely defined by cytoplasmic translation, ascospore wall assembly, mitochondrial transcription/translation, cytogamy, RNA Polymerase I initiation, protein import into the peroxisome, and DNA double-stranded break repair (full lists <u>supplementary file 2</u>). This indicates that the strongest signal coming from the yeast dataset is related to fundamental cellular processes like transcription, translation, and growth.

**Figure 5:**
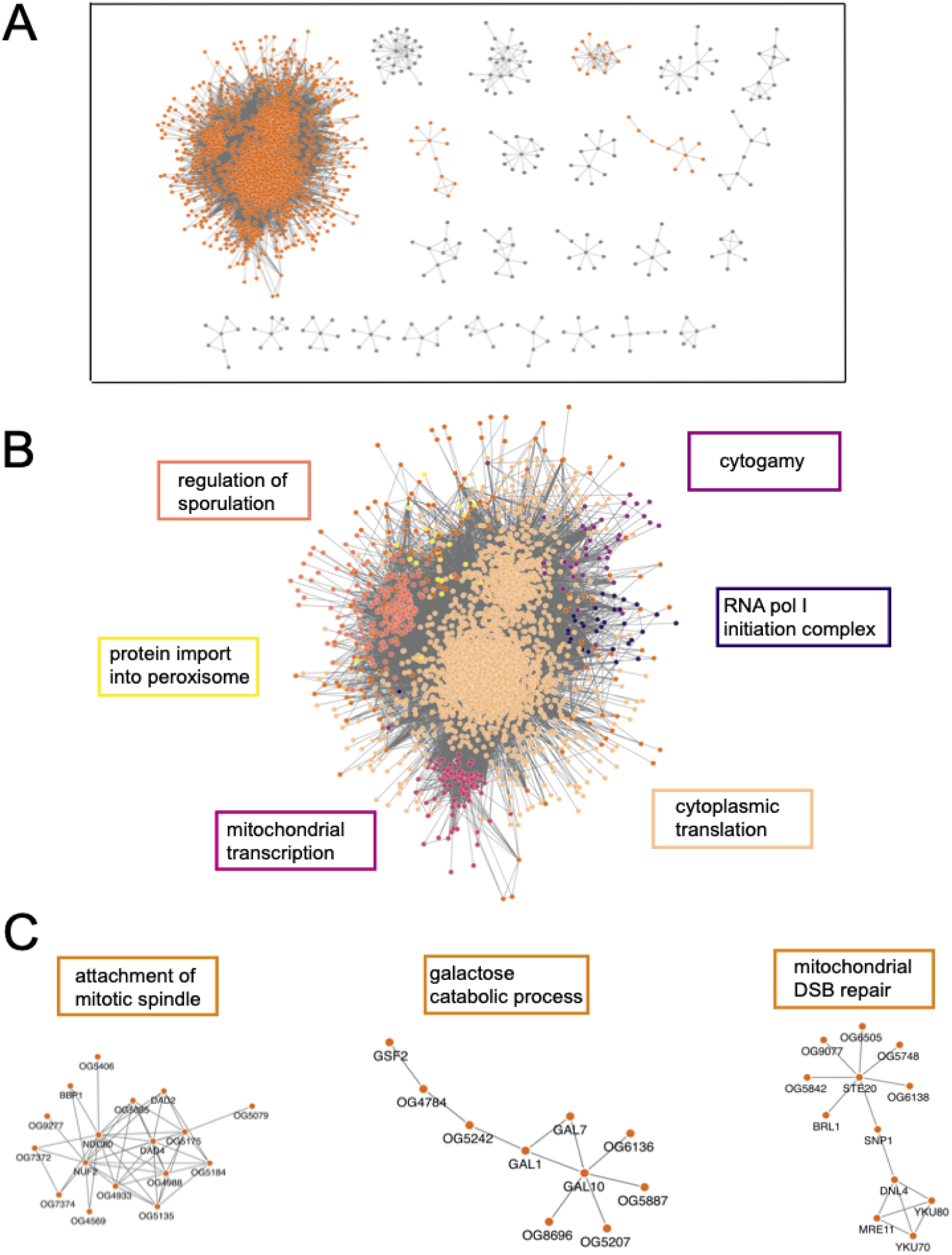
Yeast ERC is largely dominated by transcription/translation related processes. Cytoscape MCL clusters using the top 500K yeast FtERC values as the weights. A full network of all clusters is pictured in <u>Figure S4</u>. All MCL clusters with more than 6 members, clusters highlighted in B and C, are highlighted in orange. B. Enlarged image of MCL cluster 1. The large cluster is colored by subclusters with the highest enriched GO:Biological Process term listed next to the subcluster in the corresponding color. C. Select clusters with the highest enriched GO: Biological Process term is shown on top of the cluster (DSB = double-stranded break). The full enrichment lists are in supplementary file 2.

The yeast networks also make specific predictions of new genes in each of these functional categories, because each network contained some genes not previously appreciated to be in the enriched function (<u>Table S2</u>). Some of these novel genes were completely uncharacterized, while others were already annotated to unrelated functions, which suggests an additional, pleiotropic function. Other novel genes are not in the *Saccharomyces cerevisiae* genome at all, in which most functional annotation has occurred. They were instead present in many non-*cerevisiae* yeast species. Those novelties show the potential to assign functions to genes in non-model species, including in core processes shared more broadly across taxonomic groups. For example, we observed several non-*cerevisiae* genes that had high ERC values with recognized mitochondrial genes; those predictions were upheld in databases from non-*cerevisiae* yeast species (Supplementary file 7).

In contrast, the mammal dataset formed networks capturing more organismal level biological processes. The mammalian network, similar to the yeast network, has one large cluster containing 1504 of all 4805 nodes. (Figure 6, <u>Figure S5</u>, supplementary file 3). After sub-clustering, there were 3 major sub-groups of cluster 1 that were enriched for a variety of GO Biological Process terms such as brain development, limb development, and spermatid development (supplementary file 4). Among the clusters with 10 or more members, there are many organism-level enrichment terms such as stem cell fate specification, animal organ formation, and detection of chemical stimuli involved in sensory perception of smell. While there are clusters that capture transcription/translation (2, 13, and 24), overall, the mammalian network shows that ERC is capturing more organismal-level biological processes compared to the yeast network, which is heavily dominated by processes involved in transcription and translation.

**Figure 6:**
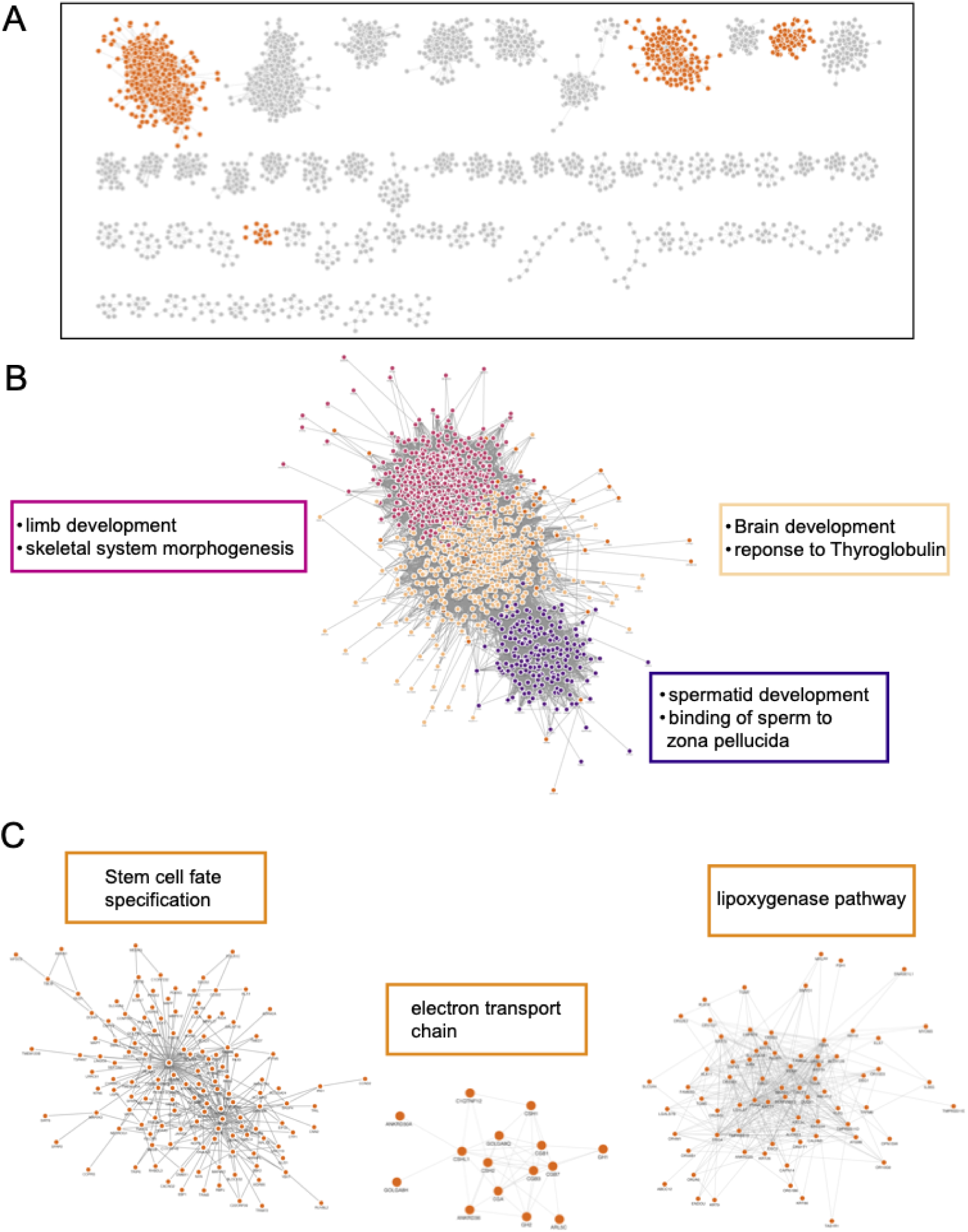
Mammal ERC captures organismal processes. Cytoscape MCL clusters using the top 500k mammal FtERC values as the weights. A. Clusters with 10 or more members, all network clusters can be found in <u>Figure S5</u>, clusters highlighted in B and C are highlighted in orange. A. B. Enlarged image of MCL cluster 1. The large cluster is colored by subclusters with the highest enriched GO:Biological Process term listed next to the subcluster in the corresponding color. C. Select clusters with the highest enriched GO: Biological Process term shown on top of the cluster. The full enrichment lists are in <u>supplementary file 3</u>.

### Case Study: ERC2.0 identifies known and novel functional interactions with histone chaperones

To demonstrate that ERC2.0 analysis is sufficiently informative to identify both known and novel relationships between genes, we examined transcription-associated histone chaperones as a test case using the 343 yeast species dataset. Histone chaperones interact with histones and modulate the disassembly and assembly of nucleosomes in an ATP-independent manner during DNA-templated processes (Hammond et al. 2017). In *Saccharomyces cerevisiae*, Spt6, Spn1, and the FACT (FAcilitates Chromatin Transcription) complex, containing Spt16 and Pob3, function as histone chaperones during transcription by RNA Polymerase II (RNAPII) (Miller et al. 2023; Robert and Jeronimo 2023). All four of these proteins are conserved in mammalian cells and are essential for the viability of yeast cells, indicating that they perform critical independent functions. Recent genetic and genomic work in yeast suggests that despite being essential individually, these factors cooperate to maintain proper chromatin architecture in the wake of RNAPII transcription (Viktorovskaya et al. 2021; López-Rivera et al. 2022; Warner et al. 2024).

To understand the functional coordination among these histone chaperones, we asked which genes have a high ERC with *SPT16*, *POB3*, *SPT6*, and *SPN1*. Since the distribution of ERC values for each gene was different (<u>Figure S6A</u>), we standardized the results by calculating Z-scores for ERC values in each distribution (<u>Figure S6B</u>). In this section, we define a gene pair to have a high ERC value if it passes a Z-score cut-off of greater than or equal to 3.00, which selects roughly the top 1% of genes with each histone chaperone. We visualized genes that have a high ERC value with the histone chaperone genes as a network (Figure 7, supplementary file 5).

**Figure 7:**
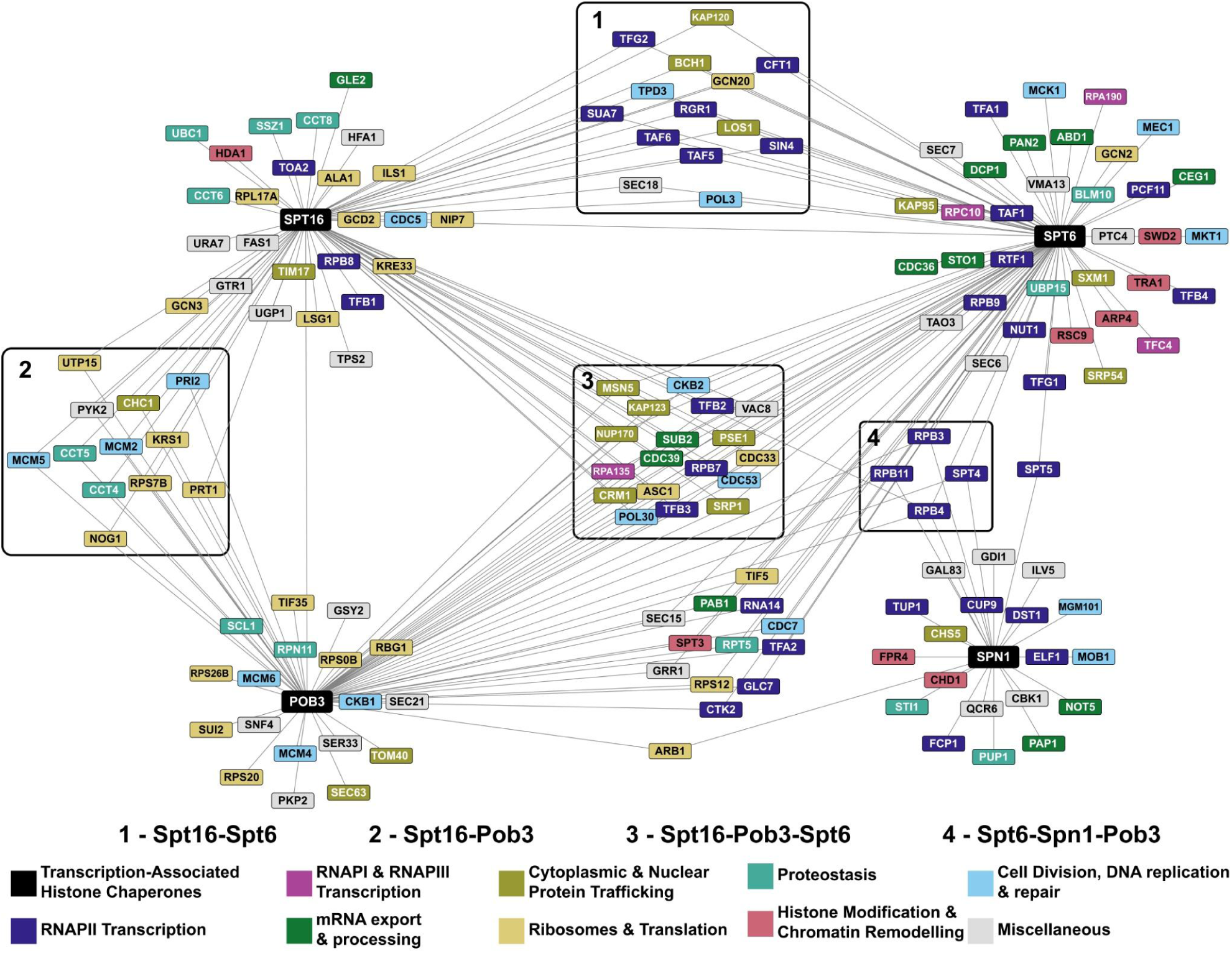
ERC network of transcription-associated histone chaperones identifies both known and putative functional interactions. Each node in the network is a gene and an edge between two nodes indicates that the genes share an ERC value above a Z-score cut-off of 3 (*SPT6*: N = 72, *POB3*: N = 67, *SPT6*: N = 86, *SPN1*: N = 25). The ERC values in this network range from 6.94 to 16.01. Clusters in boxes represent nodes that are connected to two or more of the queried histone chaperones. The colors of the nodes represent the biological process associated with the gene. Biological process associations were manually curated from the *Saccharomyces* Genome Database. The miscellaneous category consists of genes involved in autophagy (*GTR1*), budding (*CBK1* and *TAO3*), metabolism (*FAS1, GAL83, GRR1, GSY2, HFA1, ILV5, PKP2, PYK2, QCR6, SER33, SNF4, TPS2, UGP1,* and *URA7*), vesicular transport (*GDI1, SEC15, SEC18, SEC21, SEC6,* and *SEC7*), vacuole acidification (*VMA13*), and vacuole fusion (*VAC8*).

To ensure that this network was informative of the functional relationship between the genes and not random, we compared the global clustering coefficient of this network to that of 10,000 randomly generated networks. The histone chaperone network had a higher global clustering coefficient than all 10,000 sampled networks, indicating a significant degree of connectivity between the genes that shared a high ERC value with the histone chaperones (<u>Figure S6C</u>).

Several genes in the network are connected to two or more of the queried histone chaperones (Figure 7, clusters one to four). These shared genes support existing literature showing that transcription-associated histone chaperones functionally coordinate with one another (Viktorovskaya et al. 2021; López-Rivera et al. 2022; Warner et al. 2024). Genes involved in the nuclear import of proteins, RNAPII subunits, and transcription elongation factors have a high ERC value with *SPT16*, *POB3*, and *SPT6* (Figure 7, cluster three). In addition to their roles in transcription elongation, *SPT16* and *SPT6* have been implicated in transcription initiation, and expectedly, several genes involved in the process (Figure 7, blue nodes in the cluster one) have a high ERC value with both histone chaperones (Biswas et al. 2005; Doris et al. 2018). Genes encoding the DNA replication factors Pol3 and Pol30 have a high ERC value with *SPT16, POB3,* and *SPT6,* and genes encoding subunits in the Mcm2-7 complex have a high ERC value with FACT. This is consistent with Spt6, Pob3, and Spt16 having functions in DNA replication (Formosa and Winston 2020; Miller and Winston 2023; Wang et al. 2023). Thus, the ERC network indicates functional interactions between the histone chaperones and proteins central to nuclear processes impacted by chromatin structure. However, the absence of an edge between a histone chaperone and another node in this network does not imply a lack of functional connection. For example, although Spt6 physically interacts with Mcm2 and Mcm4 and has been implicated in DNA replication, an edge was not drawn between the two as the ERC values were just under the Z-Score cut-off (Z-score*_SPT6-MCM2_* = 2.88 and Z-score*_SPT6-MCM4_* = 2.95) (Supplementary File 6) (Miller and Winston 2023). We do note that Spn1, which physically interacts with Spt6 in the transcription elongation complex (Farnung 2025), has a narrower ERC value distribution (<u>Figure S6A</u>) and is less connected to other nodes in the network.

We also asked if the network suggests previously unexpected connections between transcription-associated histone chaperones and other biological processes. Interestingly, some translation and ribosome-associated genes have a high ERC value with *SPT16* and *POB3* (Figure 7, yellow nodes in cluster two). There are even broader relationships captured by the edges between the black and grey nodes. These edges suggest relationships between proteins involved in transcription-associated histone chaperoning with functions involved in metabolism and vesicular transport, among others. This may be due to direct relationships between the histone chaperones and these other functions, or individual proteins on either side of these edges may play a non-canonical role elsewhere that is contributing to the high ERC score. These observations highlight the power of ERC analysis to generate hypotheses to examine underappreciated relationships between factors in pathways thought to be disparate.

## Discussion

ERC2.0 allows for faster and more accurate computation of evolutionary rate covariation for large species datasets. We show that for a dataset of 343 yeast species or 120 mammalian species, there is an improvement in predictive power over using the *S. cerevisiae* and *H. sapiens* STRING databases alone. The new methodology provides increased power over older methods due to improvements to normalization of rates (removing heteroskedasticity) and correlation measures (Fisher transformation), which also allows better comparisons between ERC datasets calculated in different taxonomic groups. Most gained power however comes from using more species. Interestingly, as more species are added, there seems to be a plateau of power above 147 species in yeast. We attribute some of this to the fact that we are limited to annotations in *S. cerevisiae,* and some of the high-scoring pairs in the 196-343 species bins are specific to clades outside of *S. cerevisiae* and, therefore, would not be recorded as a true positive. There could also be a plateau because approximately 100 species are enough to capture the majority of variation in selective environments encountered in that taxonomic group.

ERC can be used to rank genome-wide interactions with a gene/complex/pathway of interest to identify candidate interactors and has had great success predicting previously unknown functional relationships valuable to experimental biologists (Findlay et al. 2014; Raza et al. 2019; Priedigkeit et al. 2015; Clark et al. 2013; Kowalczyk et al. 2021). We also built ERC-based functional gene networks using MCL clustering, which suggest new co-functional interactions and discovered a discrepancy in the types of processes and functions captured by the highest-scoring edges for yeast and mammals. In the yeast networks, we saw a very strong cluster of transcription/translation-related genes, whereas the mammalian network captured more macro-level terms, such as sperm motility. These differences could indicate a difference in the variation of evolutionary pressures experienced by single-celled yeast versus multicellular mammals. This pattern could also be an artifact of the much greater evolutionary distance captured in the yeast species, where the majority of overlap between orthologous genes is captured by essential processes such as transcription. The latter hypothesis will require further exploration of individual clades within the 343 yeast dataset by splitting the phylogeny into different subgroups of species, running ERC on each individually, and assessing the differences in FtERC scores for a given pair.

We introduced the application of a logistic regression model to make functional predictions which could be applied for any set of annotations either from a database or a custom annotation specific to a given study. We showed that for the 343-yeast species dataset, there is strong predictive power for two different annotation sets. There was less predictive power for the mammal ERC dataset. However, across all three datasets, 60% of the terms still performed with an AUC greater than 0.8, which indicates useful predictive power, and when employed in a predictive capacity, the user would know how well ERC performed on known genes (using ROC-AUC), before deciding whether to pursue new candidates suggested by the model.

We also showed the potential for ERC to aid researchers who work in non-model systems. First, the yeast dataset makes many strong functional predictions for non-*S. cerevisiae* genes, many of which were validated after consulting annotations from non-*cerevisiae* species (Supplementary file 7). Second, the combination of ERC with logistic regression predictions allows for a re-assessment of already annotated genes. In some cases, an experimental assay shows a gene/protein as a part of a specific function, which creates a confirmation bias whenever that gene/protein shows up in other contexts. By looking at candidate interactions in an unbiased way, such as ERC, we may uncover novel functions of proteins beyond their current functional annotations. Third, even more potential for non-model systems is demonstrated by the fact that ERC alone has predictive power approaching that of the entire STRING database (<u>Figure S3A</u>). Considering that STRING draws from experimental and computational predictions amassed over decades by thousands of researchers, having ERC as a rapid and low-cost alternative to making functional predictions will be a huge boost to non-model systems, especially because ERC only requires a collection of genomes from related species.

We also present a specific example in transcription-associated histone chaperone genes *SPT16*, *POB3*, *SPT6*, and *SPN1* to illustrate the potential of ERC2.0-derived networks as a screen for novel interactions with genes/functions of interest. The network showed edges between genes involved in related functional categories as well as suggested relationships between functions such as translation and histone chaperoning that have not yet been explored. In general, the studies presented here demonstrate how ERC2.0 could be applied in the systems biology field as a complement to other functional and interaction prediction methods. The current study was limited to yeast and mammal species, however, there are increasingly larger collections of related species genomes and more refined phylogenies, both of which promise greater power in those systems. Some more neglected taxonomic groups, such as bacteria and parasites, are of great interest to the health community, and ERC will allow researchers to rapidly identify functions and interactions within their genomes. These predictions would be especially useful in species that are more difficult to study in the lab.

## Data availability

Dryad: 10.5061/dryad.6m905qg8q

https://github.com/nclark-lab/erc

## Acknowledgments

The support and resources from the Center for High-Performance Computing at the University of Utah are gratefully acknowledged.

We acknowledge key funding from the NIH as R01 HG009299 to JHL and NLC and R35 GM141964 to KMA.

## Methods

### Generating sequence alignments and gene trees

To calculate ERC we first need to align and generate gene trees for each of the orthologous genes. For the dataset of 343 yeast, the orthologous groups (OG) were sourced from Shen *et al*. 2018. Each OG was then aligned using the MUSCLE alignment algorithm (Edgar 2004).

For the 120 mammal dataset, the coding sequence alignments were generated from the whole genome alignment found in Hecker and Hiller, 2020. First the protein-coding portions were extracted from the whole-genome alignment based on the canonical hg38 UCSC human gene models. The exons were then extracted from the MAF file using sub.msa from the RPHAST package (Hubisz et al. 2011). Next, the human reading frame was enforced and stop codons were masked as gaps using codon.clean.msa in the RPHAST package. The sequences were translated into amino acid sequences using translate.msa in RPHAST. Finally, gene trees for both sets of alignments were calculated using the phangorn (Schliep 2011), estimatePhangornTreeAll wrapper included with RERconverge (Kowalczyk et al. 2021).

### Efficient Computation of Correlations between Features

The complexity of ERC combinations stems from the fact that each feature (*e.g*., orthologous gene set) exhibits a unique presence-absence pattern across species. These patterns are largely driven by technical artifacts and evolutionary gain/loss and are effectively random, resulting in each pair of features typically sharing a unique set of species. To compare tree-encoded data between features, they must first be standardized onto a common species set by pruning the original trees. Since this operation needs to be performed separately for every pair of features, it is critical to minimize the computational cost of tree operations.

Our implementation addresses this challenge in two ways. First, because tree pruning corresponds to turning paths into edges, we precompute all possible paths resulting from any pruning operation for each tree and store them in a feature-by-path matrix. Second, we construct an indexing structure that, given a common species set, quickly retrieves the subset of paths corresponding to the tree pruned to match that set. This approach allows for a matrix indexing lookup, returning the results as a vector for fast correlation computations.

These optimizations leverage highly efficient C-based tree operations provided as helper functions in the R (R Core Team 2024) TreeTools package (Martin R. Smith 2019). As a result, our approach ensures that ∼80% of the running time is dedicated to actual correlation computations. The code implementing ERC2.0 is available at its repository on GitHub (https://github.com/nclark-lab/erc).

### Creating clustered ERC networks

To create clustered networks from ERC scores, we first subset the genome-by-genome matrices, both mammalian and yeast, to only Fisher transformed values of 10 or greater. We then imported the data into Cytoscape (Shannon et al. 2003) and clustered using the clusterMaker2 (Morris et al. 2011) MCL clustering algorithm. We kept the inflation parameter at 2.5 and removed edges between clusters for visualization purposes. The clusters were then colored by MCL cluster number along a continuous scale.

We performed GO biological process enrichment analyses on all clusters with at least 10 members for both datasets using the Gene Ontology Panther enrichment analysis tool (The Gene Ontology Consortium et al. 2023; Thomas et al. 2022) (supplementary file 2, supplementary file 4).

### Generating glmnet models and predictions

We used the glmnet (Tay et al. 2023) R package to perform 10-fold cross-validation across 2 different annotation datasets collected from Enrichr (Kuleshov et al. 2016) for each of the ERC matrices. We used the GO-BP and KEGG pathways annotation datasets.

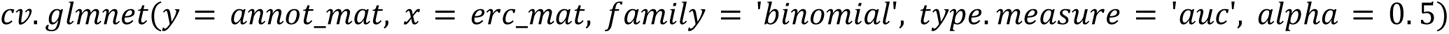

We used lambda.1se to collect the AUCs for each annotation term.

We performed two iterations of cv.glmnet for the yeast ERC matrix. The first was run with the entirety of the 12,552 orthologous genes as the input for the model. In the second matrix, only genes that are in both the ERC matrix and the STRING database were used, leaving the matrix with 4423 genes. The STRING-matched ERC matrix was also used to predict annotations for the non-*S. cerevisiae* orthologous genes (<u>Table S2</u>), which was run using the command:

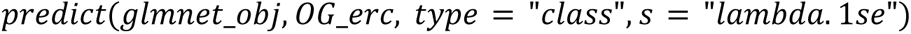

The OG_erc is a matrix of *S.cerevisiae* genes as the columns and non-*S.cerevisiae* genes as the rows. Both annotation sets were used to generate predictions.

---

**Figure S1.**
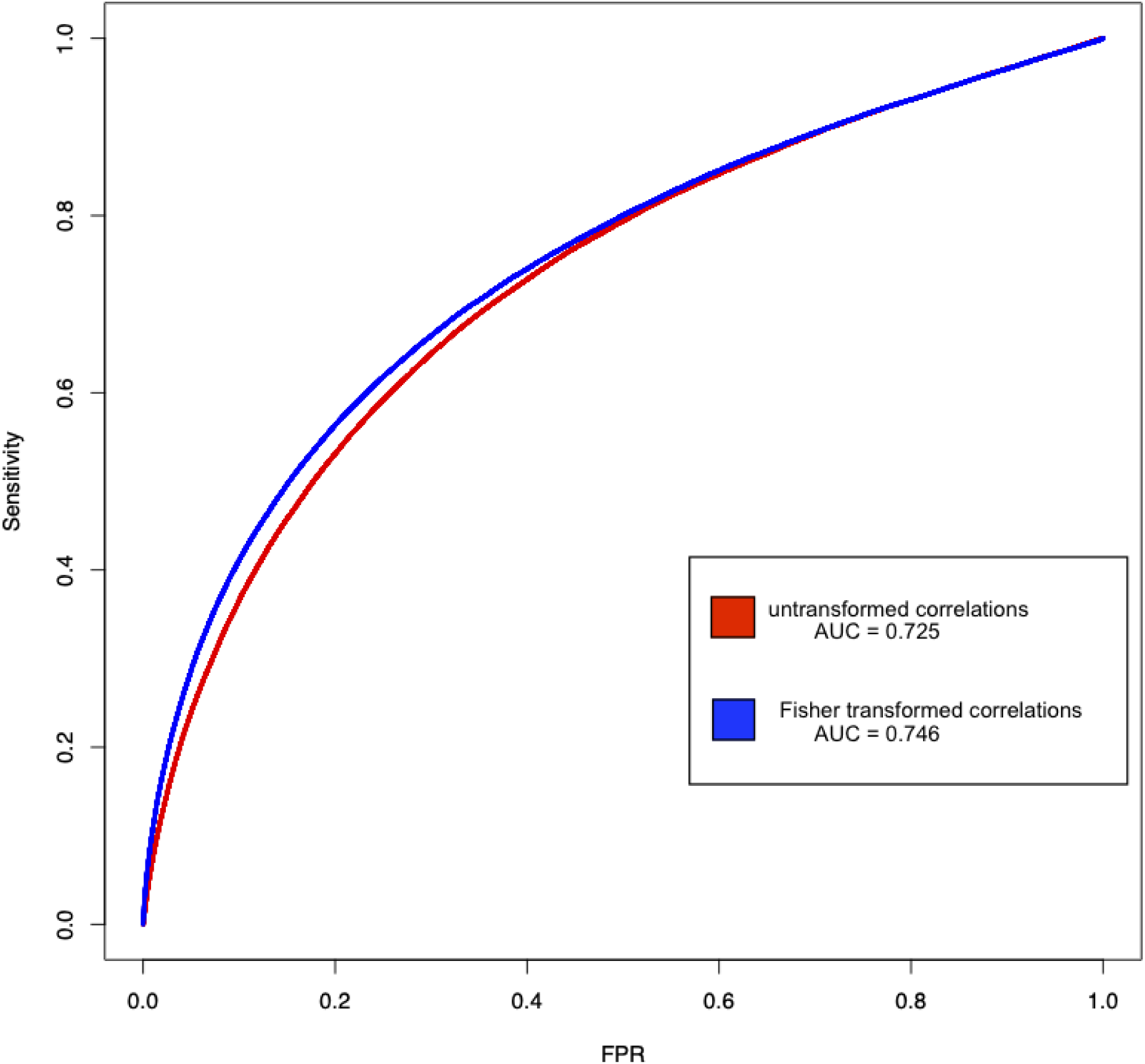
Fisher transformed correlations outperform untransformed Pearson correlations when comparing ERC to the STRING database. ROC curves comparing the predictive power of untransformed ERC correlations to Fisher transformed correlations for ERC calculated on 343 yeast when using the STRING database as the truth set. The red curve shows a ROC curve calculated on untransformed Pearson correlations and resulted in a ROC-AUC of 0.725. The blue line shows the ROC curve calculated using Fisher transformed Pearson correlations and resulted in a ROC-AUC of 0.746

**Figure S2:**
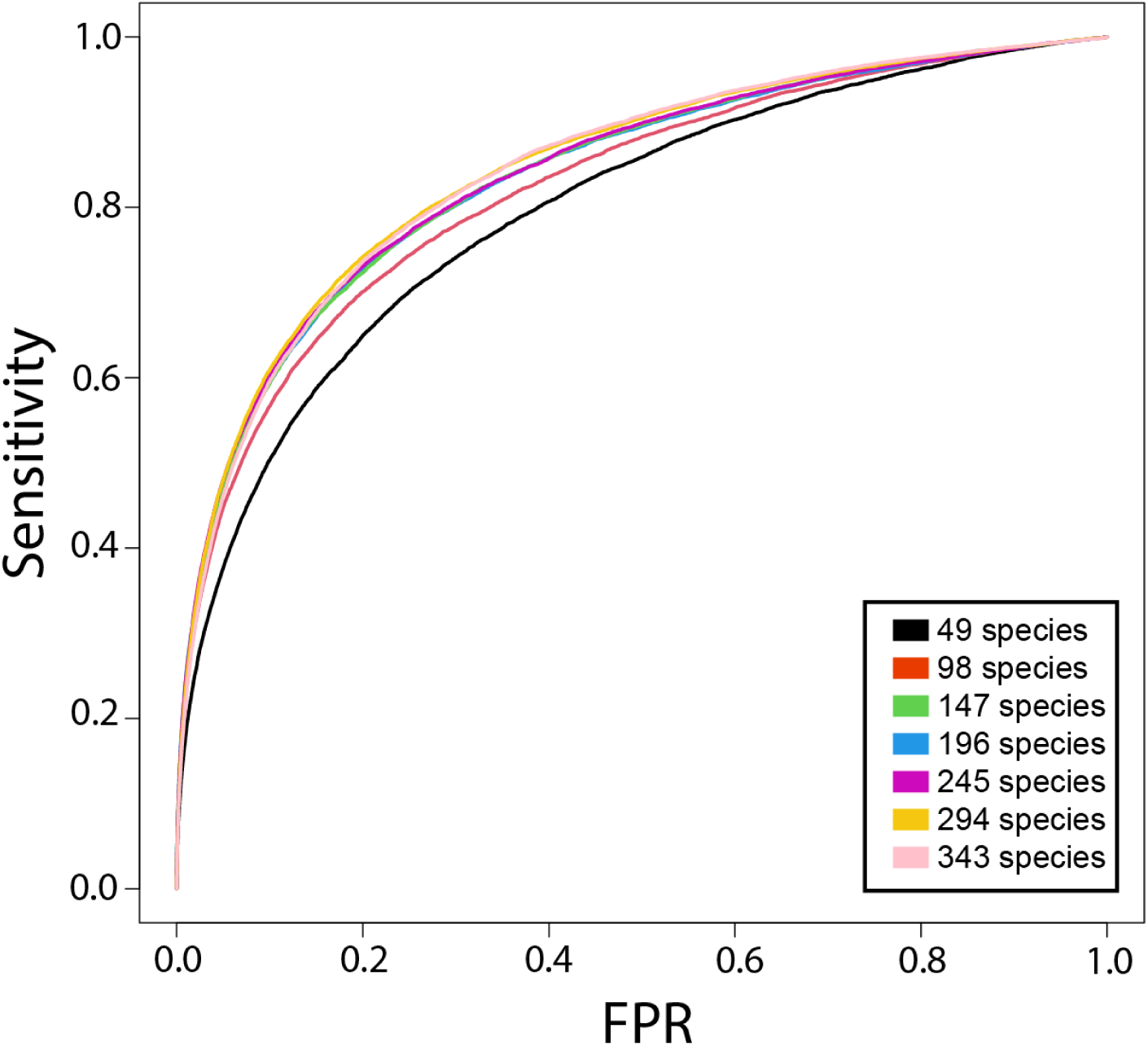
ERC2.0 power to predict STRING interactions increases with species number until a plateau at 147 species. ROC curves for increasingly larger subsets of yeast species from the whole dataset of 343 yeast species. Each subset increases by 49 species from the previous. ROC curves were calculated using STRING interactions as the truth set. FPR = false positive rate

**Figure S3:**
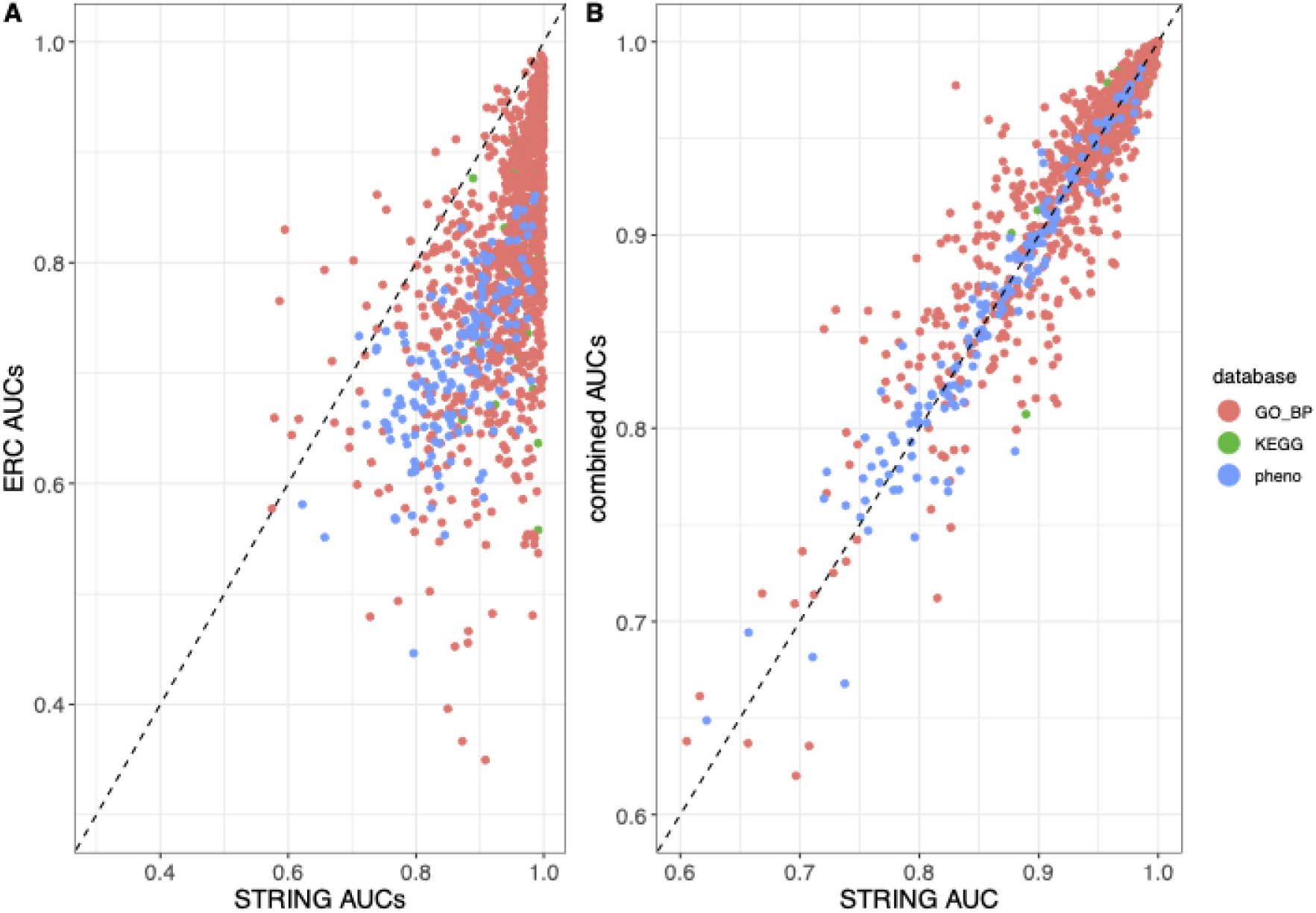
Logistic regression models generated with STRING scores outperform those generated from ERC2.0 scores in yeast. A. Scatterplot comparing the cv.glmnet lambda.1se AUC values using either STRING (x-axis) or ERC (y-axis) as the input data for three datasets in yeast: KEGG (green), Phenotypes (blue), and GO: Biological processes (red). Thirty annotation terms are above the diagonal, 29 from GO: Biological processes and 1 from phenotypes. B. Scatterplot comparing the cv.glmnet lambda.1se AUC values using either STRING (x-axis) or combined STRING/ERC (y-axis) as the input data for three datasets in yeast: KEGG (green), Phenotypes (blue), and GO: Biological processes (red).

**Figure S4:**
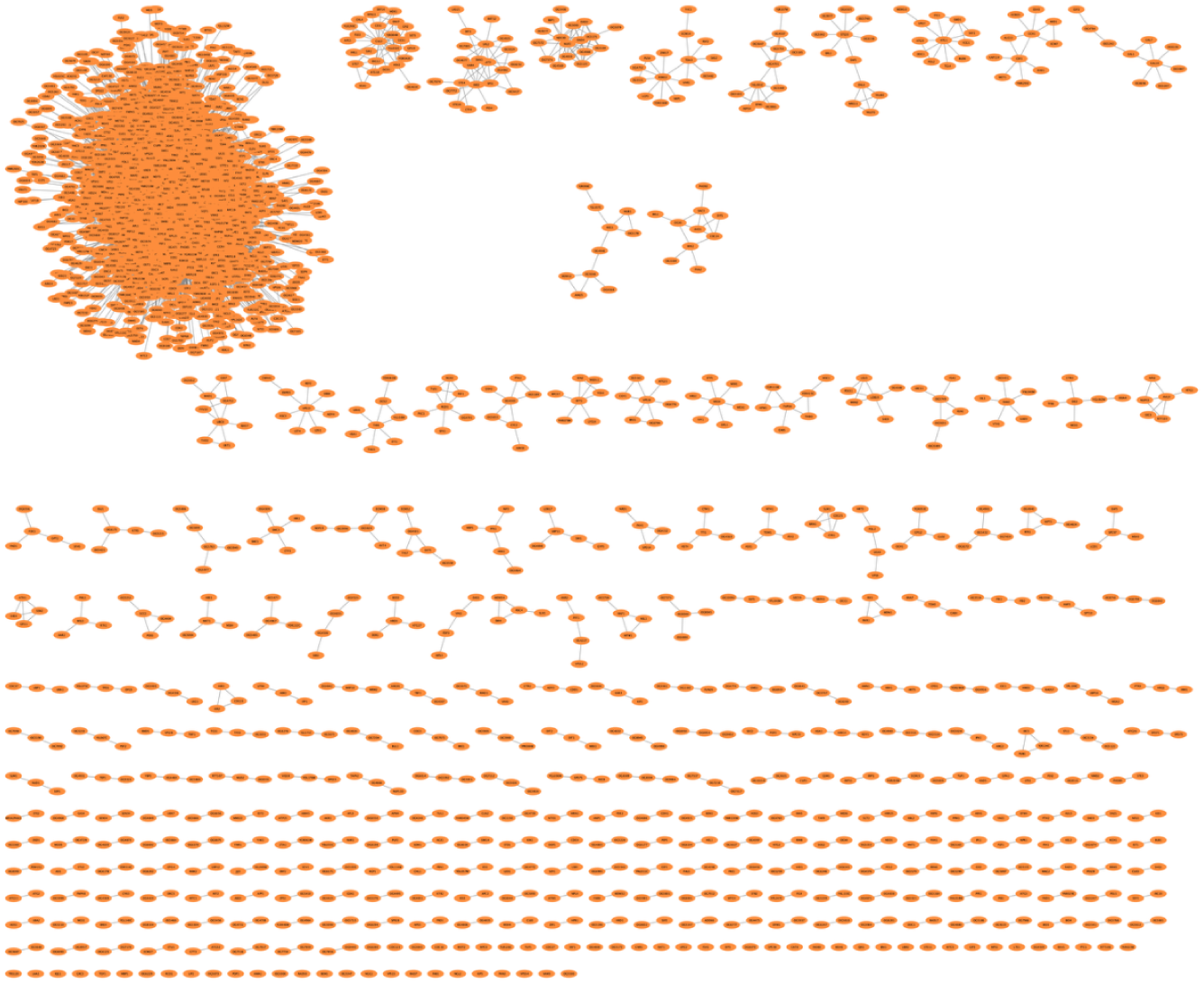
ERC2.0 edges calculated using 343 yeast species are largely group in one cluster. MCL clustering of the top 500 thousand FtERC edges of the 343 yeast dataset. Each orange node represents a gene. The lines between nodes are ERC2.0 scores. Edges between clusters were removed for easier visualization.

**Figure S5:**
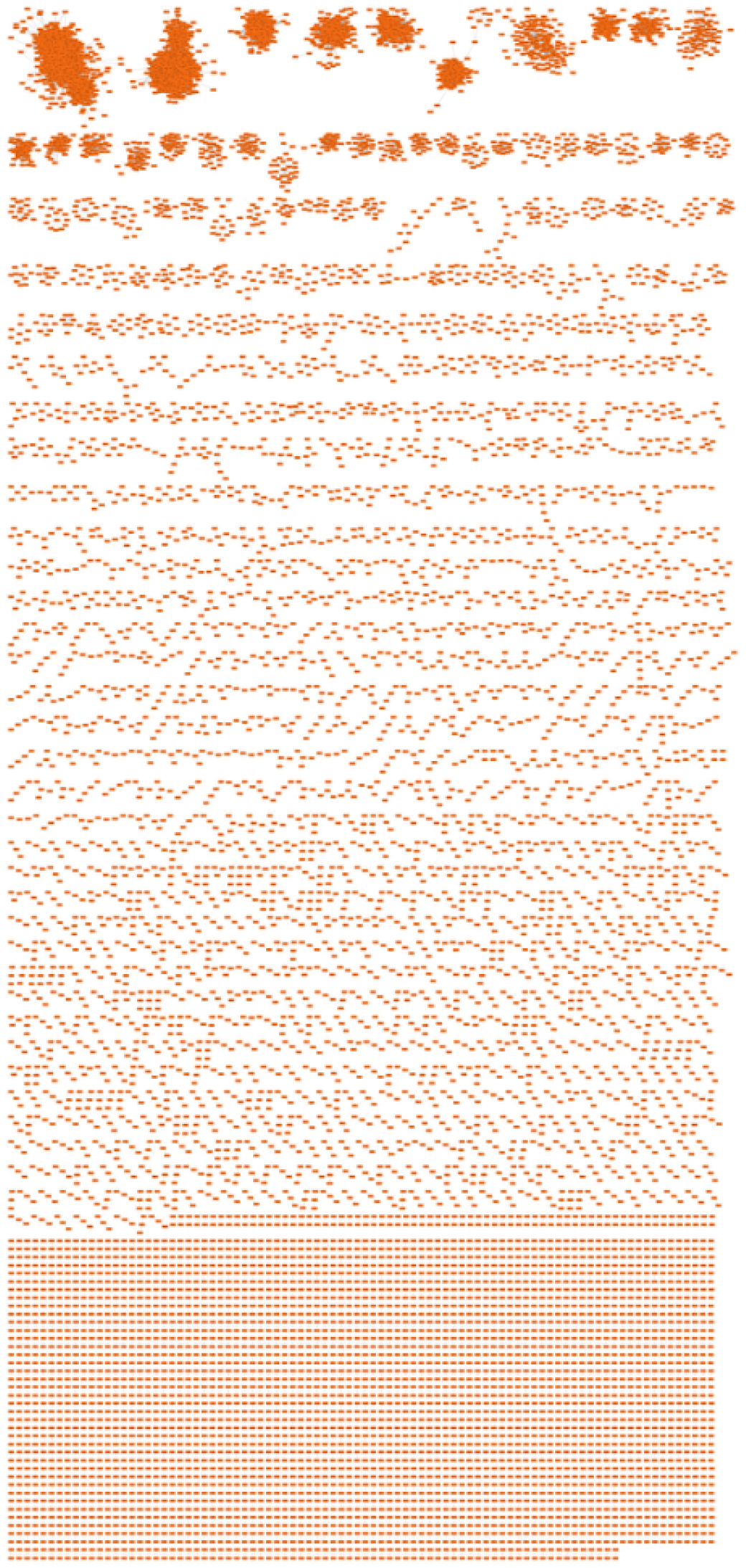
MCL clustering of the top 500K FtERC edges in the mammal dataset ERC2.0 edges calculated using 120 yeast species have many subclusters. MCL clustering of the top 500 thousand FtERC edges of the 120 mammal dataset. Each orange node represents a gene. The lines between nodes are ERC2.0 scores. Edges between clusters were removed for easier visualization.

**Figure S6:**
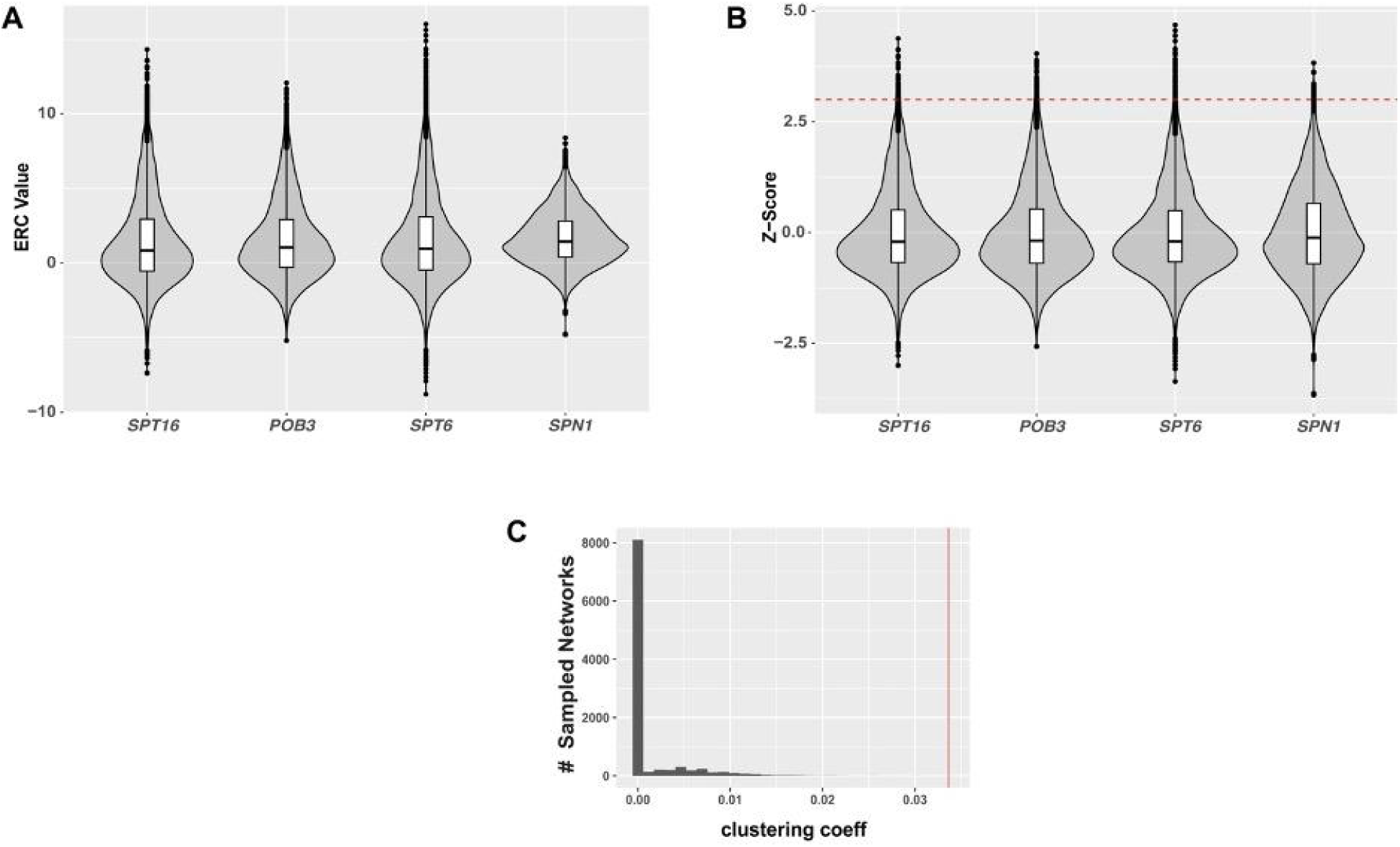
The transcription-associated histone chaperone ERC network has a higher global clustering coefficient than randomly sampled networks. A) Violin plot showing the distribution of ERC values for *SPT16*, *POB3*, *SPT6*, and *SPN1*. B) Violin plot showing the distribution of Z-score standardized ERC values for *SPT16*, *POB3*, *SPT6*, and *SPN1*. The red line indicates a Z-score of 3, which was used as the cut-off examine at gene pairs with high ERC values. C) Histogram of the distribution of global clustering coefficients within 10,000 randomly sampled networks. Each sampled network was generated first by randomly choosing 4 query genes. For each set of query genes, a network was prepared as in Figure 7 using the genes with the top N ERC values for the respective query where N = 72, 67, 86, or 25, thus producing networks with the same total number of edges per query node. Displayed is the global connectivity of each sampled graph as assessed using the global clustering coefficient. Vertical red line indicates global clustering coefficient of the chaperone network in Figure 7 (0.033).

**Table S1:**
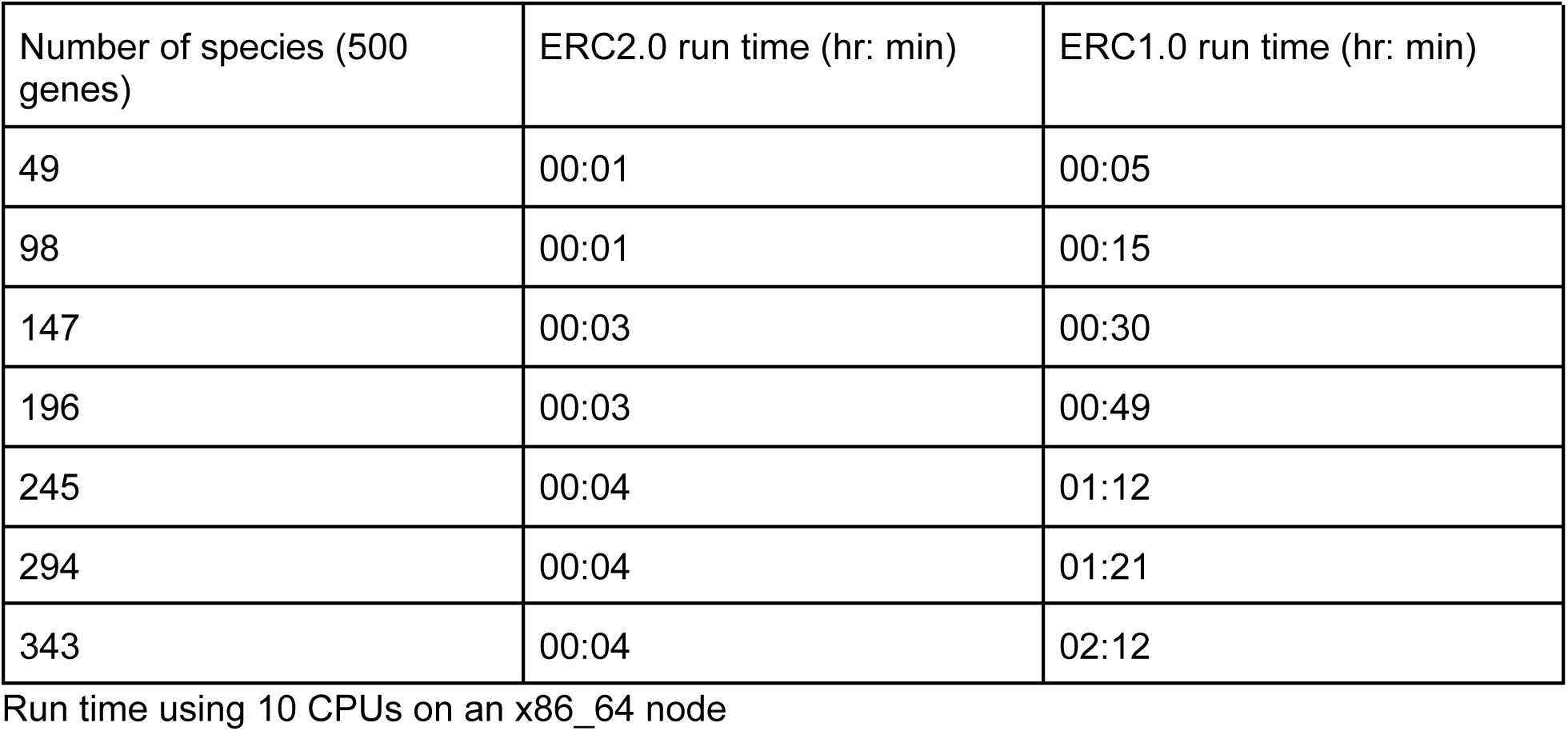

**Table S2:**

| Annotation term | Predicted member(s) |
| --- | --- |
| Oxidative phosphorylation | MME1 |
| Ribosome | OG5228 |
| Maturation of SSU-rRNA from tricistronic rRNA transcript | RPA190 |
| Ribosomal large subunit biogenesis | UTP14, BMS1, MRD1, RPA190 |
| translation | RPO41, MAM33, MBA1, YGR021W, OG4591 |
| DNA-dependent DNA replication | MEC1 |
| RNA splicing, via transesterification reactions with bulged adenosine as nucleophile | PRP8, PRP5, PRP28, OG4742 |
| Cell wall mannoprotein/glycoprotein biosynthetic process | YJL181W |
| mRNA splicing, via spliceosome | PRP8, PRP5, MUD2, PRP28, OG4742, OG5197 |
| Mitochondrial respiratory chain complex IV assembly/biogenesis | YLR283W |
| DNA-templated transcription, elongation | TAF1 |
| mRNA processing | PRP5, PRP28 |
| Respiratory electron transport chain | YMR166C |
| Pre-replicative complex assembly involved in nuclear cell cycle DNA replication | OG4420 |
| phosphorylation | OG3228, OG4757 |
| Regulation of transcription from RNA polymerase II promoter | TAF5, BDF1, UBP2, OG7669 |
| Mitotic recombination | RFA1, RFA2 |
| Septum formation | OG7034 |

